# Optimising fertilisation kinetics models for broadcast spawning corals in the genus *Acropora*

**DOI:** 10.1101/2025.05.13.653772

**Authors:** Elizabeth Buccheri, Russell C Babcock, Peter J Mumby, Christopher Doropoulos, Gerard F Ricardo

## Abstract

Synchronous spawning is a specialised adaptation to maximise fertilisation success in free spawning sessile invertebrates. There are many factors that drive and limit reproduction in such benthic invertebrates including localised hydrodynamic forcing, adult density dependence, and fine-scale gamete interactions, all of which are difficult to measure *in situ*. Therefore, measures to manage or restore reproductive populations of sessile invertebrates rely in part on adequate modelling of spawning processes at localised scales. Fertilisation kinetics of spawning events has been modelled for many free spawning marine invertebrate taxa, yet little work has been done to parameterise such models for hermaphroditic corals. This study used experimentally derived coral-specific parameters and optimisation protocols to improve model predictions of fertilisation outcomes for *Acropora kenti* (formerly *A*. “Maggie” *tenuis*) and *A. digitifera*. Three fine scale parameters that are difficult to measure experimentally – fertilisation efficiency (*Fe*), egg concentration (*E0*), and polyspermy block strength (*tb*) – were estimated using optimisation, and medians and 95% confidence intervals were derived for each parameter and species. For *A. kenti* the median optimised *Fe* value was 0.0194 (0.0027–0.0776), and *tb* was 0.1000 (0.1000–0.1000). For *A. digitifera*, the median optimised *Fe* value was 0.0013 (0.0002–0.0030), and *tb* was 10.0000 (2.4703– 10.0000). Further, sensitivity analyses suggest that kinetics models are the most sensitive to changes in *Fe* parameter values, as well as species-specific metrics like sperm swimming speed and egg size. Results will inform biophysical coral fertilisation models to better predict reproductive outcomes and support management decisions that safeguard natural reef recovery.

## Introduction

Coral reefs are facing declines in coral density and species diversity in the Anthropocene (Connell, Hughes and Wallace 1997, De’Ath et al. 2012, Emslie et al. 2024). The thinning of coral populations is expected to have direct implications for density-dependent processes that govern natural population recovery, including reproduction via mass spawning (Teo and Todd 2018, Mumby et al. 2024, Ricardo et al. 2024). For example, Allee effects can limit reproduction in the absence of adequate fecund adult densities and will affect both individual fitness and population level dynamics if severe (Oliver and Babcock 1992, Knowlton 2001, Gascoigne et al. 2009). The complexity of coral spawning and fertilisation processes, as well as their brief and episodic annual nature, makes it difficult to measure the outcomes of reproduction under various scenarios *in situ*. Therefore, developing robust models to predict spawning outcomes is integral to not only understand how disturbances may impede natural recovery, but also promote management which safeguards reproductive mechanisms (McLeod et al. 2019).

Fertilisation presents particular challenges for free spawning marine invertebrates, especially those with limited mobility (Dulvy, Sadovy and Reynolds 2003). Low population densities, rapid dilution of sperm in seawater and limited gamete viability periods mean that ensuring gametes experience contact at high enough concentrations and long enough periods of time for sufficient fertilisation requires particular adaptions (Gribben, Millar and Jeffs 2014). For all benthic invertebrates, successful fertilisation relies on many interacting parameters. At fine scales, sperm concentration and egg-sperm contact time are the most influential (Levitan, Sewell and Chia 1991, Buccheri et al. 2023). These parameters can be mechanistically modelled to predict fertilisation outcomes given their relationships with other relevant parameters such as sperm swimming speed (Morita et al. 2006) and egg size (Levitan 2006). Most of the foundational reproductive literature is focused on mobile invertebrates like sea urchins and abalone (Vogel et al. 1982, Levitan, Sewell and Chia 1991, Styan 1998, Babcock and Keesing 1999), with little focus on corals despite their importance as reef-building and sessile external fertilisers.

The most prominent fertilisation kinetics model for spawning individuals, named Vogel-Czihak-Chang-Wolf (VCCW) was initially created by Vogel et al. (1982) using the sea urchin *Paracentrotus lividus*. This model was derived from biomolecular kinetics and developed in response to early simulations conducted by Rothschild and Swann (1951) and Hultin and Hagstrom (1956). The VCCW model presented fertilisation success as a function of egg size, egg concentration, fertilisation efficiency, sperm concentration, sperm half-life, and sperm swimming velocity. The model assumed that the fertilisation site, where the sperm attaches to the egg and the proportion of successful fertilisation encounters, correspond to egg size (Vogel et al. 1982, Jantzen, de Nys and Havenhand 2001). The VCCW has had iterative improvements (known as the extended VCCW) to include polyspermy block components that further refine predictions of fertilisation success (Styan 1998, Millar and Anderson 2003). This model has been widely referred to and adapted for numerous free-spawning invertebrates (Levitan 1993, Jantzen, de Nys and Havenhand 2001, Lehtonen and Dardare 2019).

Fertilisation success typically increases non-linearly with sperm concentration, following a sigmoidal trend (Levitan, Sewell and Chia 1991, Benzie and Dixon 1994, Franke, Babcock and Styan 2002, Levitan and McGovern 2005). Fertilisation rates are generally low when sperm concentrations are low, due to few egg-sperm collisions (Franke, Babcock and Styan 2002), and these increase exponentially as sperm concentration increases (Pennington 1985, Levitan and Petersen 1995, Baird, Guest and Willis 2009). In some species, as sperm concentration exceeds an upper threshold, successful fertilisation rates may decline, likely owing to the influence of low dissolved oxygen (DO) (Oliver and Babcock 1992, Simpson, Cary and Masini 1993) or polyspermy (Styan 1998, Millar and Anderson 2003). Polyspermy occurs when an egg is fertilised by multiple sperm, and can result in embryo abnormalities or death (Brawley 1987, Gribben, Millar and Jeffs 2014). The degree of influence of polyspermy on coral reproduction is species-specific because some genera, like most acroporids (Morita et al. 2006), have eggs with efficient polyspermy blocks that activate following contact with sperm to prevent the negative impacts of over-exposure at biologically relevant sperm concentrations (Styan 1998). Thus, lack of dissolved oxygen may be a more likely limiting factor for *Acropora* at high sperm concentrations.

Contact time between sperm and eggs is also an important factor that influences fertilisation (Buccheri et al. 2023), but the degree of influence is species-specific (Nozawa, Isomura and Fukami 2015). For example, Nowaza et al. (2015) found that fertilisation rate increased as contact time increased from 10 to 30 to 60 minutes for four of their study species, *A. gemmifera, Favites abdita, F. pentagona*, and *F. valensiennesi*, but had no influence on fertilisation rates for *A. papillare* and *Platygyra ryukyuensis* (Nozawa, Isomura and Fukami 2015). While Buccheri et al. (2023) observed more distinct positive trends in fertilisation success as contact time was increased from 10 seconds to 30 minutes incrementally for *A. kenti, A. digitifera, P. daedalea, and Coelastrea aspera*. Following spawning, corals have a specific amount to time to mix and fertilise before they become too diluted to interact (Omori et al. 2001). Therefore, contact time is a crucial parameter, and is dependent on the density of fecund colonies in an area, water depth, and the severity of hydrodynamic mixing experienced to mix the spawn (Denny and Shibata 1989, Nozawa, Isomura and Fukami 2015). For example, if wind-driven hydrodynamic forcing is strong, sperm can quickly get diluted making fertilisation unlikely, especially in deeper water (Wolanski, Burrage and King 1989, van Woesik 2010).

Gamete properties drive fine-scale interactions that may promote or inhibit fertilisation (Levitan, TerHorst and Fogarty 2007). Sperm swimming characteristics influence gamete contact at small scales (Farley 2002), often due to the presence of chemoattractants in eggs (Yoshida, Inara and Morisawa 1993, Coll et al. 1994, Morita et al. 2006). For example, it has been observed that sperm from many species swim faster, and change directional patterns, in the presence of conspecific eggs (Morita et al. 2006). Sperm swimming speed is variable across taxa (EB, RCB, CD, GFR personal obs), and chemoattraction capabilities are species-specific (Evans and Sherman 2013) and rapidly evolving (Metz and Palumbi 1996, Levitan and Ferrell 2006), implying varied influences on fertilisation success. Egg size has also been observed to promote successful fertilisation, but there is a trade-off between egg size and fecundity which species likely evolved independently (Levitan 2006). Egg concentration is not likely a significant driver of fertilisation success (Lillie 1915, Levitan, Sewell and Chia 1991), thus is expected to have little influence on fertilisation kinetics. Species-specific gamete properties result in variability in reproductive capacity, thus require consideration when modelling fertilisation outcomes.

Many foundational concepts underpinning reproductive biology are consistent across spawning invertebrates. However, the fine-scale nuances of fertilisation kinetics are variable across groups due to vast differences in spawning tactics, gamete properties, and resulting hydrodynamic exposure during spawning (Babcock 1995, Crimaldi and Zimmer 2014). Thus, fertilisation kinetics models need to be parameterised for specific systems, and species of interest, to improve the accuracy of predictions. The present study aimed to validate and optimise kinetics models for two species of hermaphroditic spawning corals in the Indo-Pacific: *A*. cf. *kenti* and *A*. cf. *digitifera*, using coral-specific metrics. Model accuracy has improved greatly for the study species, and results are expected to inform biophysical coral spawning models. More realistic modelled outcomes will improve understanding of species-specific demographic processes, ultimately informing more effective management decision-making.

## Methods

### Egg size estimation for *A. digitifera*

Egg sizes were measured for *A. digitifera* during the 2017 spawning event at Coral Bay Research Station in Western Australia. Five colonies of *A. digitifera* were collected from a reef flat in Coral Bay (23.1423° S, 113.7723° E) and brought to the laboratory prior to the full moon of their anticipated spawning month. Gravid colonies were selected after snapping a branch and observing pigmented eggs. Corals were kept in ambient free-flowing aquaria and were isolated into still 40L containers and observed for setting (the emergence of egg-sperm bundles to the polyp mouth) and spawning on predicted spawning nights (Baird et al. 2021). All colonies spawned at 21:45–22:15 on the 22nd of March, and ∼5 egg-sperm bundles were collected from each individual. Bundles were placed in 1 mL Eppendorf tubes and gently mixed to separate egg and sperm components, then eggs were rinsed to remove sperm. The total number of eggs in each bundle was documented, and each egg was measured while suspended in seawater using an optical micrometre to determine their maximum diameter (µm). Average egg sizes for *A. kenti* were gathered from the literature (Wallace 1999, Bridge et al. 2023).

### Sperm swimming speed estimation for *A. kenti*

Sperm swimming speed assays were conducted across two nights during the 2023 spawning event at the Australian Institute of Marine Science (AIMS) National Sea Simulator facility (SeaSim). Eight *A. kenti (*formerly *A. “*Maggie*” tenuis)* colonies were collected from reefs surrounding Magnetic Island (−19.1385° S, 146.8339° E) in the central GBR and brought to the laboratory prior to the full moon of their anticipated spawning month. Colony selection and spawning protocol followed the same methods outlined in the previous section. Four colonies spawned on 2 November at 18:15-18:40, and three colonies spawned on 3 November at 18:15-18:25. Roughly 100 bundles were collected from each colony and kept isolated. For each colony, egg-sperm bundles were broken apart, sperm were separated from eggs using a 125-µm mesh sieve, and eggs were rinsed with filtered seawater (FSW) and left to incubate in 10 mL of 1.0-µm FSW immediately for ∼20 minutes while sperm samples were processed, allowing potential release of chemoattractants by eggs. Sperm was evenly pooled from respective donors, and sperm counts were conducted using a haemocytometer to quantify concentration.

Sperm samples were diluted using either clean FSW or egg-soaked FSW to a concentration of 10^5^ sperm mL^-1^ for analysis. Sperm samples were transferred to plain glass slides with coverslips and examined at 40x magnification with phase contrast using a <u>**Zeiss Primostar 3**</u> trinocular compound light microscope with a <u>**CMOS 8.3 MP camera attachment**</u>. Sample analysis was randomised across treatments to account for any changes in sperm viability over time. All analyses were conducted within the ideal sperm viability window of 4 hours post-spawning (Willis et al. 1997). Videos of sperm were taken using the ImageView software (YSCTechnologies 2024) at 60 frames per second (FPS) for 3 seconds. Three videos were taken per slide at three different locations around the slide, and this was repeated five times for each replicate cross.

Sperm swimming was analysed with a point tracking method using the DLTdv8a app (Hedrick 2008) in Matlab version 23.2 (r2023b) (MathWorks 2023). Initial (*v*_i_) and final (*v*_f_) velocity parameters were used to calculate the average velocity (*v*_avg_) of all motile sperm in each video. Some videos were omitted due to poor quality or lack of visible or motile sperm for long enough durations (>2 sec). A total of 82 total sperm tracks were quantified across 35 videos. Sperm swimming speeds for *A. digitifera* were extracted from Morita et al. (2006) using PlotDigitizer (PlotDigitizer 2024).

Sperm swimming speed was analysed as a function of egg presence to determine whether exposure to egg-conditioned FSW contributed to faster sperm swimming speeds. Sperm swimming speed data was not-normally distributed, thus a Wilcoxon rank-sum test was used from the *stats* package in RStudio 2024.04.0 (RStudio 2024) to compare sperm-swimming speeds across the two treatment groups. Cliff’s delta was also calculated to confirm the minimal effect size of the comparison using the *effsize* package (Torchiano 2020). All analyses were conducted in RStudio using R version 4.3.2 (RCoreTeam 2023).

### Model fitting and parameter optimisation

Model parameters for coral-specific gamete properties were derived from laboratory observations for each species where possible, otherwise from the literature (Table 1, Fig. 1). Additional parameters like the cross-sectional area of an egg (*σ*) and the collision rate constant (*β*) were calculated within the model using the known parameters from the literature and this study. The cross-sectional area of an egg was calculated using *σ*= *πr*^2^, where r is the radius of an egg in mm. The collision rate constant is calculated with *β* = ν σ, where ν is sperm swimming speed (mm s^-1^) and σ is the cross-sectional area of the egg (mm^2^).

**Table 1.** Model parameters with known values based on literature or experimental data

| Species | Biological parameters | Value | Unit | Source |
| --- | --- | --- | --- | --- |
| *A. kenti* | Sperm swimming speed ($v$) | 0.19 | $\text{mm s}^{-1}$ | This study |
| | Egg diameter ($\sigma$) | 0.550 | mm | Wallace (1999)<br>Bridge et al. (2023) |
| | Sperm concentration ($SO$) | $10^0 - 10^5$ | $\text{mL}^{-1}$ | Buccheri et al. (2023) |
| | Contact time ($time$) | 10, 30, 60, 600, 1800 | s | |
| *A. digitifera* | Sperm swimming speed ($v$) | 0.30 | $\text{mm s}^{-1}$ | Morita et al. (2006) |
| | Egg diameter ($\sigma$) | 0.588 | mm | This study |
| | Sperm concentration ($SO$) | $10^0 - 10^5$ | $\text{mL}^{-1}$ | Buccheri et al. (2023) |
| | Contact time ($time$) | 10, 30, 60, 600, 1800 | s | |

**Fig. 1.**
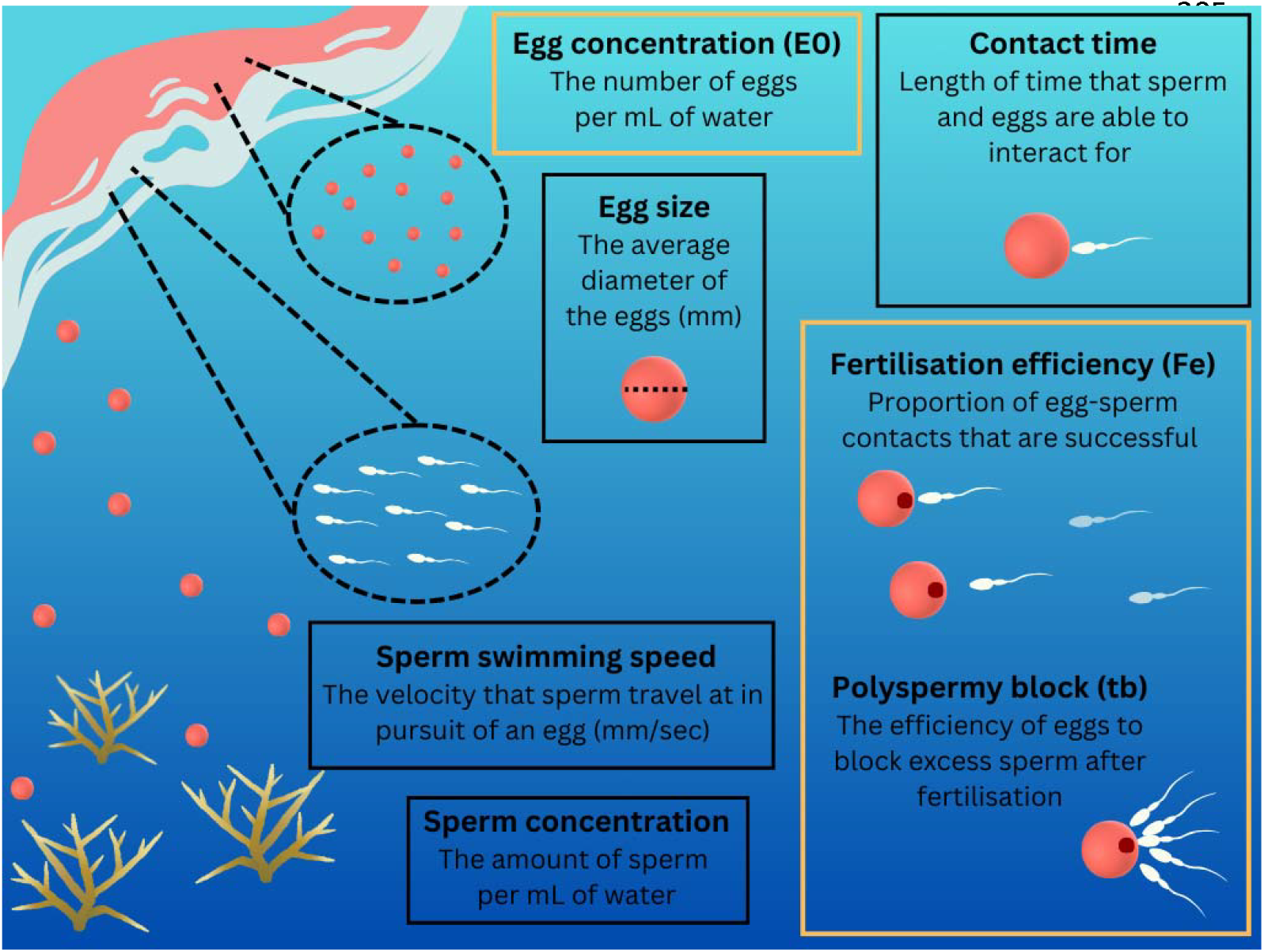
Schematic figure depicting the spawning process and outlining the areas that fertilisation kinetics modelling focuses. Parameters outlined in black were known based on published literature or experimentation, and parameters outlined in yellow are predicted using optimisation protocols

The Extended VCCW (EVCCW hereafter) model was used to optimise the remaining model parameters: fertilisation efficiency (*Fe*), egg concentration (*E0*), and polyspermy block time (*tb*).

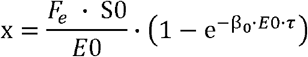

where *x* is the average number of potential fertilisers per egg

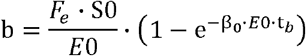

where *b* is the mean number of extra fertilising sperm that will contact an egg in the time *tb*, of eggs already contacted once.

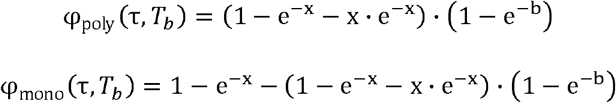

Model fits were compared to observed results from laboratory trials for each species (Buccheri et al. 2023). Briefly, previous fertilisation trials were conducted by exposing ∼100 eggs to conspecific sperm at varying concentrations and for incremental contact times from 10 seconds to 30 minutes. Fertilisation success was quantified across treatments for *A. kenti* and *A. digitifera* respectively. Preliminary analyses of model fit were conducted to determine adequate ranges of each parameter to test, using expected values from the literature.

*Fe*, or fertilisation efficiency, is the fraction of egg-sperm contacts that are potentially fertilising (Styan 1998). Few estimates of *Fe* have been calculated in the literature for other spawning invertebrates, with values ranging from 0.00137 to 0.0512 (Denny and Shibata 1989, Levitan, Sewell and Chia 1991, Styan and Butler 2000, Millar and Anderson 2003). Corals are expected to have similarly low *Fe* values, but this has never been explicitly quantified.

In the EVCCW model, the egg concentration *E0*, is expected to absorb a proportion of the sperm, leading to fewer egg-sperm collisions per egg at higher egg concentrations, especially above 0.01 eggs µL^-1^ (Fig. S1). Typical ranges of egg concentrations *in situ* are unknown; however, Ricardo et al. (2024) found that for *A. kenti*, there was no evidence of egg concentration impacting fertilisation success from 2 to 300 eggs mL^-1^ (0.002 – 0.3 eggs µL^-1^). As *E0* used in this study was 0.002 eggs µL^-1^ (Buccheri et al. 2023) and below the range tested, we cautiously allowed *E0* to vary within a conservative range of 0.001 to 0.005 eggs µL^-1^ to allow for uncertainty in this parameter at lower values.

Lastly, the polyspermy block time, *tb*, has been quantified for other spawning invertebrates (Styan 1998), but is expected to be shorter for *Acropora* corals due to their efficient polyspermy blocks (Morita et al. 2006). Lower *tb* values signify quicker activation of polyspermy blocks (Morita et al. 2006), and less detrimental impacts on fertilisation success at high sperm concentrations (Styan and Butler 2000).

Once the likely ranges of each parameter were established for each species (Table S1), the fertilisation kinetics model was run using *nlsLM* from the nlsLM2 package (Grothendieck 2024), within a bootstrapped model fitting procedure using the boot package (Canty and Ripley 2024) to determine the combination of values for each of the three parameters that resulted in the best fits. A precise set of starting values were tested within each parameter range and all combinations of the three parameters were assessed, totalling 3,375 combinations, to account for high sensitivity to starting value fluctuation in kinetics models. The best fit model with a set of optimised parameters was selected if it had the lowest residual sum of squares, meaning it explained the most variation in the model.

Following optimisation, bootstrapping was conducted to run 200 trials with replacement for each contact time respectively to estimate uncertainty for each parameter. Maximum likelihood parameter optimisation and visualisation of model predictions compared to observed results from laboratory trials were used to determine the values of each of the three parameters that resulted in the best fits. Each contact time was analysed separately to allow for any changes in parameter estimates over time following spawning. Median parameter estimates and 95% confidence intervals for each species and contact time were calculated using the outputs.

Sensitivity analyses were conducted to evaluate the influence of changes of each parameter across its likely range on fertilisation success predicted by the kinetics model. Upper and lower values for *Fe, E0*, and *tb* were selected based on bootstrap model outcomes, and observed fluctuations in egg size and sperm swimming speed were determined based on laboratory observations to examine relevant ranges of each parameter (Table 2). Correlations between *Fe* and *tb* were assessed to determine if these parameters were independent. Where feasible, the *cor* function from the stats package in RStudio (RCoreTeam 2023) was used to quantify the Spearman correlation between the two parameters.

**Table 2.** Parameter ranges used in the sensitivity analysis based on values reported in the literature or measured experimentally (see Table 1). These values cover ∼95% of the data variability, based on the optimisation outcomes for Fe, E0, tb, and based on observations and published literature for sperm swimming speed and egg size

| Species | Parameter | Bounds |
| --- | --- | --- |
| *A. kenti* | Fe | lower = 0.001, upper = 0.100 |
| | E0 ($\text{mL}^{-1}$) | lower = 0.001, upper = 0.005 |
|  | tb | lower = 0.050, upper = 0.150 |
| | Sperm swimming speed ($\text{mm s}^{-1}$) | lower = 0.150, upper = 0.250 |
|  | Egg size (diameter, mm) | lower = 0.300, upper = 0.700 |
| *A. digitifera* | Fe | lower = 0.0003, upper = 0.003 |
|  | EO (mL<sup>-1</sup>) | lower = 0.001, upper = 0.005 |
|  | tb | lower = 2.000, upper = 10.000 |
|  | Sperm swimming speed (mm s<sup>-1</sup>) | lower = 0.200, upper = 0.400 |
|  | Egg size (diameter, mm) | lower = 0.500, upper = 0.700 |

### Water chemistry at elevated sperm concentrations

High sperm concentrations can affect water chemistry and subsequently inhibit fertilisation, but the degree of its influence has not been explicitly tested. An experiment was conducted with *A*. cf. *spathulata* during the December 2022 spawning event on Heron Island, southern Great Barrier Reef to establish explicit thresholds for the influence of dissolved oxygen (DO) on fertilisation success at elevated sperm concentrations.

Gravid colonies of *A. spathulata* were collected from the reef slope near Coral Gardens, Heron Reef (23.4486° S, 151.9144° E). Following spawning, sperm were separated from eggs using a 250-µm filter. Sperm from nine individuals were pooled and a series of 10-fold dilutions were created ranging from 1.5 × 10^2^ to 1.5 × 10^7^ sperm mL^-1^ and DO levels (percent saturation) were measured between 1 and 3 hours post spawning. The method for measuring DO involved stirring of a measurement probe in each sample. We note that in a small sample such as those used here, agitation can increase DO levels and therefore, resulting thresholds should be viewed as maximum value. The pH of sperm suspensions was also measured at ∼2 hrs post-spawning. The *drm* function in the drc package in RStudio (Ritz et al. 2015) was used to fit a non-linear regression to assess DO as a function of sperm concentration over time.

## Results

### Egg size estimation for *A. digitifera*

A total of 106 eggs were counted and measured across 5 individuals and 19 egg-sperm bundles. There was a mean of 6 eggs per bundle, with a minimum of 3 and a maximum of 10 observed. Egg diameter was 588⍰± ⍰ 351µm (mean⍰± ⍰ SD.; Table S2).

### Sperm swimming speed estimation for *A. kenti*

Sperm swimming speeds were compared across treatments to determine whether egg-conditioned water resulted in heightened sperm velocities due to chemoattraction (Jantzen, de Nys and Havenhand 2001, Riffell, Krug and Zimmer 2002). No statistical differences were observed between treatments, with negligible Cliff’s Delta effect sizes (p = 0.49, cliff’s delta = 0.095). Cliff’s Delta was used because it is robust to uneven sample size (Cliff 1993, Marfo and Okyere 2019), which we had (egg treatment = 26, non-egg = 56) due to omission of subpar sperm swimming videos from the analysis. The pooled sperm swimming speed for *A. kenti* had a median of 196.5 µm sec^−1^ and the 95^th^ percentile was 308.3 µm sec^-1^ (Fig. 2). Egg conditioned sperm had a median swimming speed of 188.9 µm sec^−1^ and a 95^th^ percentile of 270.2 µm sec^−1^, while sperm with no egg conditioning had a median swimming speed of 198.1 µm sec^−1^ and a 95^th^ percentile of 309.5 µm sec^−1^ (Table S3).

**Fig. 2.**
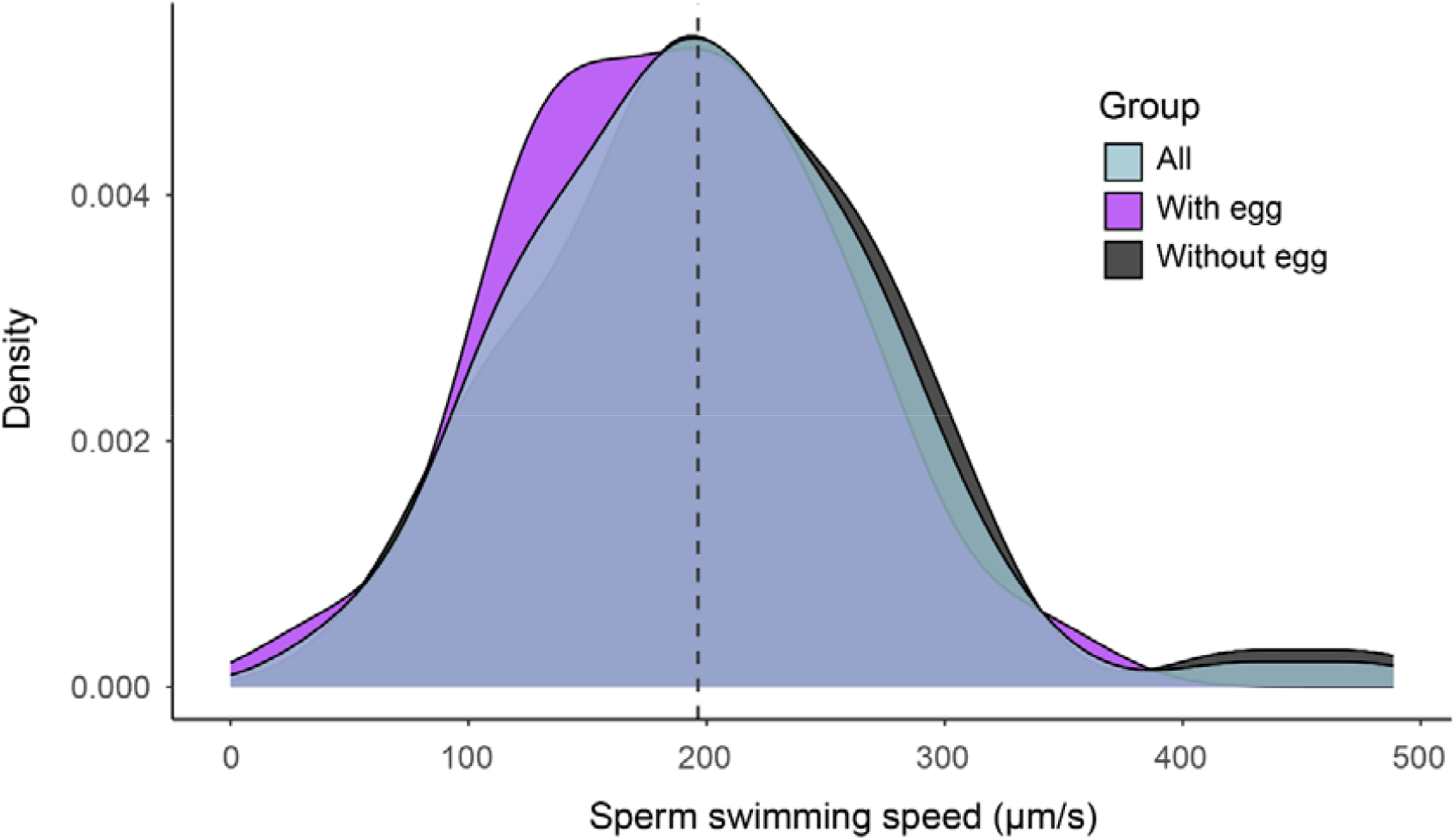
Density distribution of all sperm-swimming speed (µm s^-1^) estimates across 82 video assays for *Acropora kenti*. The black density plot represents sperm not conditioned with eggs, the purple density plot represents sperm conditioned with eggs, and the blue represents the pooled data across both groups. The dashed line represents the median of the pooled data (196.5 µm s^-1^)

### Model fitting and parameter optimisation

Predicted responses calculated using the optimised parameters initially over-or under-predicted fertilisation outcomes compared to observed data. Therefore, multiple model iterations across a range of starting values were required to produce reliable prediction estimates that supported the results from the published literature (Buccheri et al. 2023). Model fitting varied across contact times, resulting in different sets of optimised parameter values at each biological timepoint to fit the respective curves (Table S4, Fig. 3). Final optimised parameter values were summarised across all contact times for reporting. For *A. kenti* median ± 95% CI of the *Fe* parameter was 0.0194 (0.0027– 0.0776), and the *tb* parameter 0.1000 (0.1000–0.1000) (Table 3.3). For *A. digitifera, Fe* was estimated as 0.0013 (0.0002–0.0030) and *tb* was 10 (2.4703–10.0000) (Table 3).

**Table 3.** Median and 95% confidence interval distribution of optimised parameters based on 200 high precision bootstrap iterations across all contact times

| Species | Parameter | Median | Lower CI | Upper CI |
| --- | --- | --- | --- | --- |
| *A. kenti* | Fe | 0.0194 | 0.0027 | 0.0776 |
|  | E0 | 0.0050 | 0.0050 | 0.0050 |
|  | tb | 0.1000 | 0.1000 | 0.1000 |
| *A. digitifera* | Fe | 0.0013 | 0.0002 | 0.0030 |
|  | E0 | 0.0050 | 0.0010 | 0.0050 |
|  | tb | 10 | 2.4703 | 10.0000 |

**Fig. 3.**
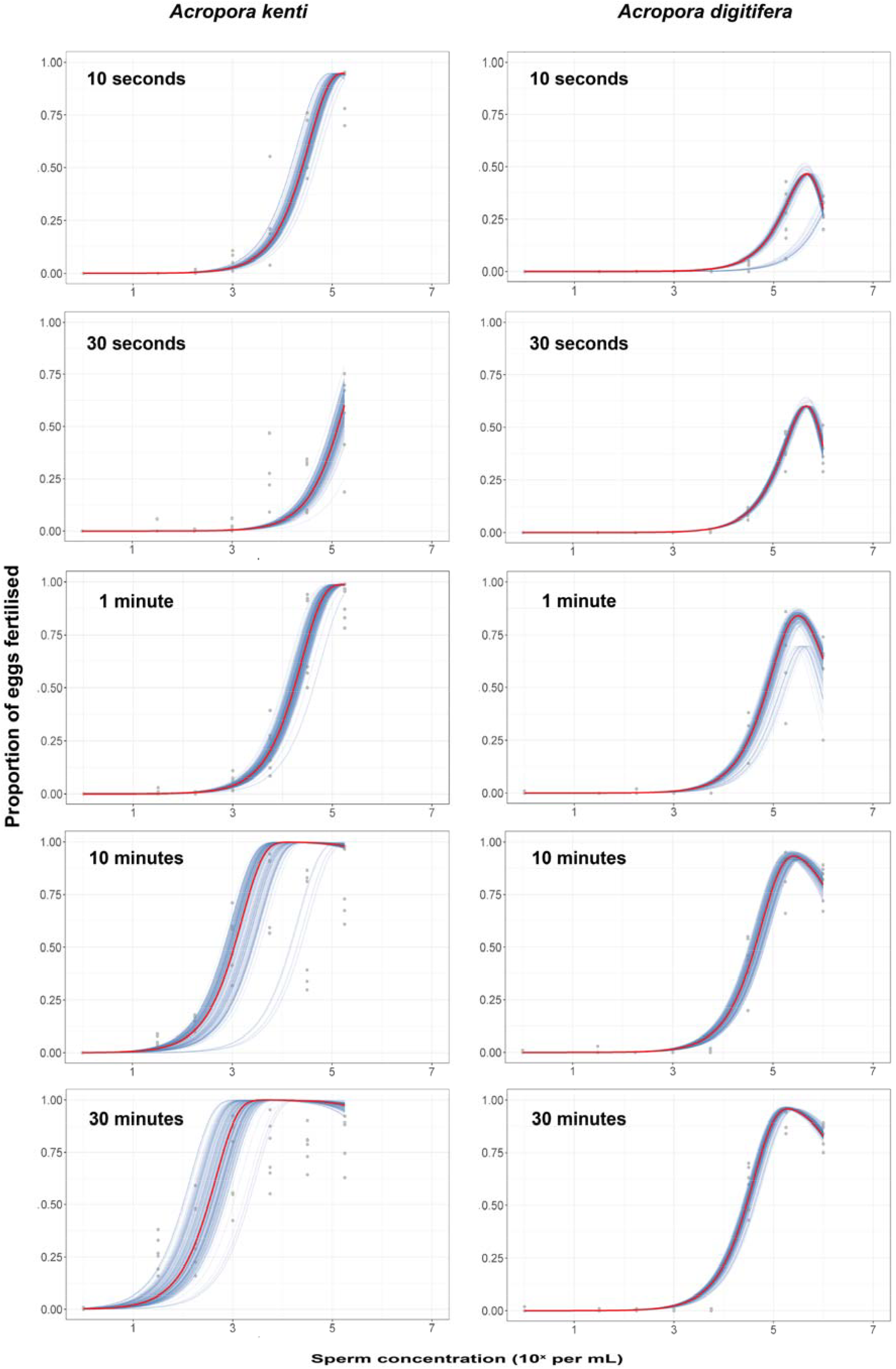
Fertilisation success as a function of sperm concentration across contact times. Points represent the raw data from fertilisation trials described in Buccheri et al. (2023). Blue lines represent fertilisation kinetics model predictions following parameter optimisation for 200 bootstrap iterations. Red lines represent the median model fits across the 200 bootstrap iterations. Each column delineates a species and panels represent each gamete contact time

Results from the sensitivity analyses conducted for both species showed that predictions of fertilisation success are highly sensitive to manipulations of *Fe* (Fig. 4). Fertilisation success was lowest, ∼5%, when the lower extreme of *Fe*’s range was tested, and highest, <95%, when *Fe* was at its upper extreme. Fertilisation success was not sensitive to changes to *E0* and *tb* within their 95% confidence ranges (Fig. 4), noting that estimates of each parameter were well known and tightly constrained within the optimisation. The fertilisation kinetics model was also sensitive to changes to sperm swimming speed and egg size within conspecific ranges, with fertilisation success ranging from ∼60–80% and 30–85% respectively (Fig. 4). Results from the Spearman correlation analysis show that there were variable levels of correlation between the *Fe* and *tb* parameters across species and contact times, ranging from minor (−0.0052) to strong (−0.7214) relationships (Fig. S2, Table S5).

**Fig. 4.**
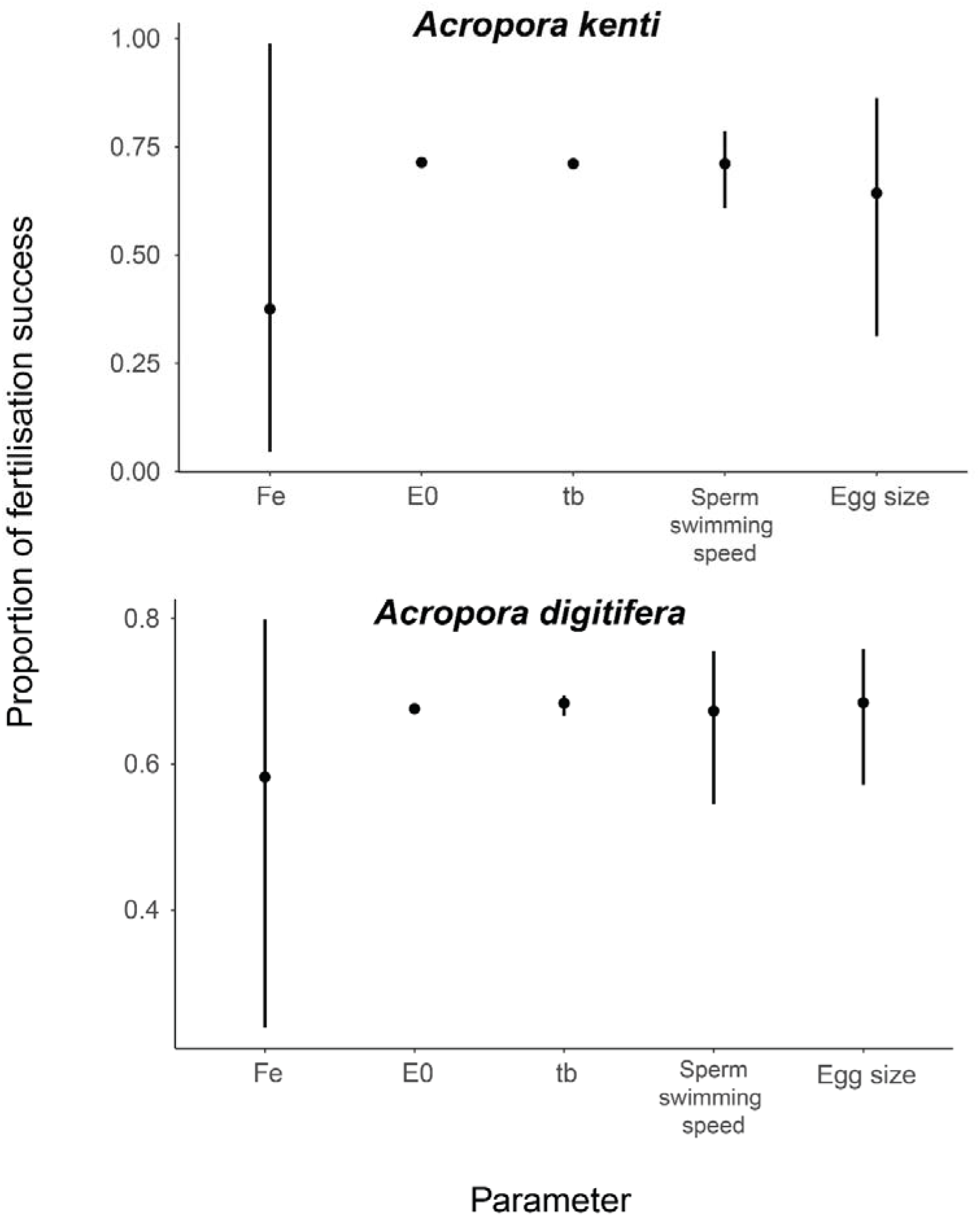
Sensitivity of predictions of fertilisation success to changes in each parameter value in the fertilisation kinetics model. Lines represent the minimum and maximum fertilisation outcomes across each parameters range. Dots represent a relative mid-range fertilisation outcome. Parameters with no line present experienced no variability in fertilisation estimates

## Discussion

The majority of fertilisation kinetics modelling in the literature has focused on other spawning invertebrates like sea urchins (Vogel et al. 1982, Styan and Butler 2000) or used values from these systems to predict outcomes for corals (Levitan et al. 2004, Teo and Todd 2018). Yet, coral spawning is unique due to differences in gamete properties and hydrodynamic processes and requires specific approaches to more accurately predict fertilisation outcomes. Further, corals have species-specific spawning behaviours (Baird et al. 2021) and gamete properties (Álvarez-Noriega et al. 2016) that further differentiate their reproductive potential. Development of specialised kinetics modelling approaches will support more comprehensive predictions of coral reproduction and build capacity for decision-making in reef management that safeguards fundamental reef recovery processes.

The fertilisation kinetics model was most sensitive to changes in the fertilisation efficiency (*Fe*) parameter, the fraction of egg-sperm interactions that are potentially fertilising (Styan 1998). This implies that the estimation of *Fe* should be a priority in any future modelling protocol to reduce uncertainty. A number of factors may reduce *Fe*, including gamete viability, egg-sperm compatibility, as well as presence of receptor sites on eggs. While the current EVCCW model only allows for *Fe* as a constant, it is likely that *Fe* varies as gametes age. Therefore, there may be some limitations in estimating a single value of *Fe*, possibly explaining some differences in *Fe* between the two species. Fertilisation kinetics model outputs were also sensitive to changes in egg size and sperm swimming speed, which highlights the importance of using species-specific metrics to reduce uncertainty.

The model was insensitive to changes in *E0*, likely because *E0* was constrained more tightly based on experimental data. Fertilisation outcomes were also insensitive to *tb*, but this will be species-specific due to variability in polyspermy block strength (Oliver and Babcock 1992, Morita et al. 2006, Lam et al. 2015). Here, *A. kenti* was tightly constrained near zero because of the little evidence of polyspermy inhibition shown in other studies (Albright et al. 2016). *Acropora digitifera* had more substantial tb, and polyspermy effects, which are unusual for acroporids. While the data indicate significant polyspermy (Buccheri et al. 2023), the estimation of *tb* is partly limited by an absence of data at high gamete concentrations above 10^7^ sperm mL^-1^. There is potential that the inhibition of fertilisation success observed could be a result of low dissolved oxygen at high sperm concentrations (Oliver and Babcock 1992, Simpson, Cary and Masini 1993), as observed with *A. spathulata* in this study (Fig. S3). We acknowledge that these values may vary slightly across species, but genus-level estimates still help contextualise this component of fertilisation kinetics. Future work should examine *A. digitifera* at high sperm concentrations to disentangle such effects.

Fertilisation kinetics outcomes are dependent on changes in sperm concentration and contact time, and these trends have been examined empirically for corals (Nozawa, Isomura and Fukami 2015, dela Cruz and Harrison 2020, Buccheri et al. 2023). The relationships are species-specific, likely due to differences in fine-scale parameters that have been shown to govern gamete interactions in other species like egg size and sperm swimming speed (Levitan 1993, Marshall, Styan and Keough 2000, Levitan, TerHorst and Fogarty 2007). Levitan (1993) found that there was a trade-off between egg size and fecundity on fertilisation success: larger eggs had higher fertilisation rates, but, as egg size increased, fecundity decreased. Therefore, organisms aim to evolve optimal egg sizes that promote fertilisation without sacrificing substantial declines in fecundity (Jantzen, de Nys and Havenhand 2001).

Such trade-offs may explain why acroporids like *A. kenti* and *A. digitifera* in this study generally have higher fertilisation rates than merulinids, that have smaller eggs, at equivalent sperm concentrations (Álvarez-Noriega et al. 2016, Madin et al. 2016, Buccheri et al. 2023). However, the relationship is often more nuanced and may also depend on gamete recognition capabilities. Chemoattractants are extremely effective at promoting fertilisation in conditions with lower than anticipated sperm-egg encounters (Jantzen, de Nys and Havenhand 2001, Levitan and Ferrell 2006). *Acropora kenti* has slightly smaller eggs and slower sperm than *A. digitifera* but had higher fertilisation success at low sperm concentrations (10^2^ – 10^4^ sperm mL^-1^). This fertilisation proficiency has been confirmed in the literature (Albright and Mason 2013, Ricardo et al. 2015) despite suboptimal gamete traits compared to other acroporids. Similar congeneric variation in fertilisation success was observed across three species of sea cucumber (Levitan 1993), and was a result of different gamete recognition capabilities at low sperm concentrations (Levitan et al. 2004). The present study did not support potential influence of chemoattractants on sperm velocity, but this may be due to the assumption that chemoattractants would exude into FSW after short exposure times (∼20 min) and would be present without physical egg-sperm contact. There is also evidence that swimming patterns vary as sperm come into contact with eggs, which also affects fertilisation (Farley 2002, Morita et al. 2006), but we had limited capacity to address this in the present study. Future studies are required to understand the drivers of fertilisation competence of *A. kenti* and discern whether chemoattractants govern their efficiency at low sperm concentrations.

Following the development and improvement of fertilisation kinetics models, as reviewed by Lehtonen and Dardare (2019), there have been major advances to predictions of reproductive outcomes for many spawning invertebrates including sand dollars, sea urchins, limpets, and more (Hodgson et al. 2007, Leuchtenberger et al. 2022). In many cases, the kinetics model overpredicted fertilisation success compared to observed experimental results, thus requiring model manipulations or additions to achieve adequate fits. In this study, optimisations regularly hit biologically informed model constraints, indicating an absence of a parameter in the model. The difficulty of fitting reproductive models to experimental data is driven by the sensitivity of kinetics models to minor changes, as well as potential biological parameters that are not explicitly accounted for. For example, Farley (2002) found that the helical nature of sperm swimming often inflated sperm swimming speed and *Fe* values and affected the fit of fertilisation kinetics models to empirical data. Thus, incorporating more detailed measurements of sperm swimming speed may enhance model performance, though these data are difficult to obtain. Here, we found the EVCCW model to be particularly sensitive to starting values during the nonlinear (weighted) least-squares optimisation. Different best fit combinations were derived from the optimisation depending on which starting values were used, therefore, we caution the use of single starting values during optimisation procedures. Further, some analyses resulted in correlations between *Fe* and *tb*. As such, optimised parameters reported here should generally be implemented together when adopted for downstream models.

The EVCCW has occasionally been used to explain coral spawning outcomes. Levitan et al. (2004) conducted laboratory and field assays to examine the mechanisms of reproductive isolation in *Orbicella* (previously *Montastraea*) *annularis* species complex and used the EVCCW to analyse their results and fit 95% confidence intervals to their data. Teo and Todd (2018) implemented the EVCCW model to examine fertilisation kinetics for corals within a simulated spawning event, using the coral *Merulina ampliata*. This spatially explicit simulation model manipulated intercolonial distance and coral colony densities to determine the implications of *in situ* adult distributions on fertilisation success. They found that fertilisation increased as colony density increased, and that larger colonies minimised the chances of sperm dilution due to their increased gamete output and concentrations (Teo and Todd 2018). A subsequent coral spawning model by Ricardo et al. (2024) likewise implemented the EVCCW and validated outputs using *in situ* spawning experiments but highlighted that some kinetics parameters such as *Fe* generated high levels of uncertainty in the modelled outcomes. Coral spawning models in the literature have drawn similar general conclusions to those based on field studies of Allee effects in spawning coral populations (Oliver and Babcock 1992, Coma and Lasker 1997, Levitan et al. 2004, Mumby et al. 2024). However, the present study demonstrates the importance of incorporating species-specific modifications to refine model predictions and improve understanding of localised coral reef contexts.

Coral reef conservation has become increasingly focused on restoration efforts like coral out-planting to safeguard reproductive populations (Bayraktarov et al. 2019). For spawning corals, efficient out-planting strategies must take reproductive potential into account to ensure spawning success and functional reef reproduction. Improved fertilisation kinetics modelling will inform appropriate population densities that safeguard natural reproduction and resilience, thus, will build capacity of reef practitioners to manage and restore local spawning coral populations.

## Compliance and ethical standards

The authors declare no conflicts of interest and confirm that all international, national, and institutional guidelines for sampling and experimentation were followed with all necessary approvals to work on the Great Barrier Reef (GBRMPA Permit G21/44774.1).

## Data statement

The datasets and code from the experimental component of this study are deposited in the CSIRO Data Access Portal http://hdl.handle.net/102.100.100/705861?index=1 and will be made fully available following publication.

## Author contributions

All authors contributed to the study conceptualisation and design. Material preparation, data collection and analysis were performed by Elizabeth Buccheri and Gerard Ricardo. The first draft of the manuscript was written by Elizabeth Buccheri and all authors provided revisions on previous versions of the manuscript. Funding was attained by Peter Mumby and Christopher Doropoulos. All authors read and approved the final manuscript.

## Acknowledgements

The authors would like to acknowledge the Traditional Owners of the Great Barrier Reef, particularly the Byelle, Gooreng Gooreng, Gurang and Taribelang Bunda First Nations people and the Wulgurukaba and Bindal First Nations people for permission to collect corals from their Sea Country and bring them to laboratory facilities for experimentation. We would like to thank Maxwell Steven, Cathy Liptrot, Tina Howe, and ZEISS Microscopy for laboratory support, Rosanna Griffith-Mumby and Anthea Donovan for logistical support, and the staff at the National Sea Simulator facility and Heron Island Research Station for assistance. We would also like to thank the reviewers for their feedback during the editorial process.

This work was supported by the EcoRRAP subprogram (https://gbrrestoration.org/program/ecorrap/) that is part of the Reef Restoration and Adaptation Program (https://gbrrestoration.org/). The Reef Restoration and Adaptation Program is funded by the partnership between the Australian Government’s Reef Trust and the Great Barrier Reef Foundation. The funders had no role in study design, data collection and analysis, decision to publish, or preparation of the manuscript.

