## Supplementary materials for "Optimising fertilisation kinetics models for broadcast spawning corals in the genus *Acropora*"

### Supplementary material

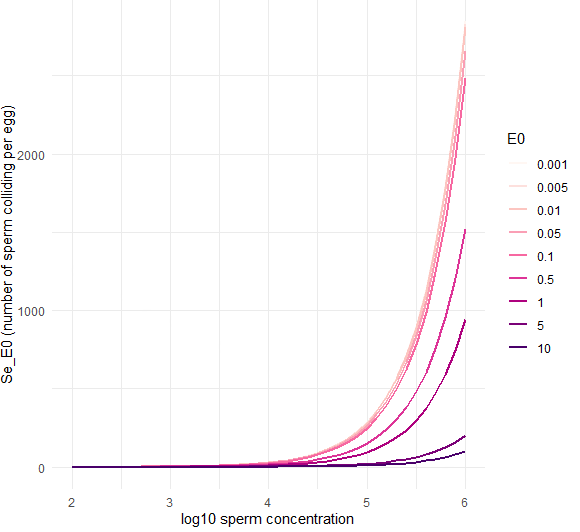

**Fig. S1** The influence of sperm concentration on the egg-sperm collisions per egg according to the EVCCW model. This simulation was run for *A. kenti* at a contact time of 60 seconds

**Table S1** Range of values tested for each parameter during the bootstrapping optimisation process for *A. kenti* and *A. digitifera*.

| **Species** | **Parameter** | **Minimum value** | **Maximum value** |
| --- | --- | --- | --- |
| *A. kenti* | Fe | 0.0010 | 0.1000 |
|  | E0 | 0.0010 | 0.0050 |
|  | tb | 0.0000 | 0.1000 |
| *A. digitifera* | Fe | 0.0001 | 0.1000 |
|  | E0 | 0.0010 | 0.0050 |
|  | tb | 0.0000 | 10.000 |

**Table S2** Egg counts and average area and diameter measurements per bundle for *A. digitifera.* Mean egg diameter was 588 with a standard deviation of 35.16.

| **Bundle** | **Number of eggs** | **Mean egg area (um^2^)** | **Mean egg diameter (um)** |
| --- | --- | --- | --- |
| 1 | 8 | 228062 | 602 |
| 2 | 6 | 226342 | 601 |
| 4 | 4 | 260000 | 609 |
| 5 | 4 | 267339 | 609 |
| 6 | 6 | 214549 | 563 |
| 7 | 6 | 227508 | 586 |
| 8 | 4 | 237117 | 583 |
| 9 | 3 | 221273 | 554 |
| 10 | 6 | 230428 | 576 |
| 11 | 3 | 230233 | 580 |
| 12 | 5 | 192722 | 514 |
| 13 | 6 | 203989 | 573 |
| 14 | 7 | 232992 | 608 |
| 15 | 5 | 232979 | 599 |
| 16 | 4 | 235945 | 609 |
| 17 | 5 | 212431 | 550 |
| 18 | 10 | 195389 | 544 |
| 19 | 6 | 309102 | 685 |
| 20 | 8 | 240898 | 626 |

**Table S3** Median and 95^th^ percentile for each sperm swimming speed group, with egg conditioning, without egg conditioning, and pooled for *A. kenti*.

| **Group** | **Median** | **95^th^ percentile** |
| --- | --- | --- |
| Egg conditioned | 188.9 | 270.2 |
| Not egg conditioned | 198.1 | 309.5 |
| Pooled | 196.5 | 308.3 |

**Table S4** Median and 95% confidence interval distribution of optimised parameters based on 200 high precision bootstrap iterations for each contact time respectively.

| **Species** | **Contact time (seconds)** | **Parameter** | **Median** | **Lower CI** | **Upper CI** |
| --- | --- | --- | --- | --- | --- |
| *A. kenti* | 10 | Fe | 0.0592 | 0.0433 | 0.1000 |
|  | 10 | E0 | 0.0050 | 0.0019 | 0.0050 |
|  | 10 | tb | 0.1000 | 0.1000 | 0.1000 |
|  | 30 | Fe | 0.0037 | 0.0023 | 0.0051 |
|  | 30 | E0 | 0.0050 | 0.0044 | 0.0050 |
|  | 30 | tb | 0.1000 | 0.1000 | 0.1000 |
|  | 60 | Fe | 0.0143 | 0.0104 | 0.0207 |
|  | 60 | E0 | 0.0050 | 0.0050 | 0.0050 |
|  | 60 | tb | 0.1000 | 0.1000 | 0.1000 |
|  | 600 | Fe | 0.0240 | 0.0018 | 0.0379 |
|  | 600 | E0 | 0.0050 | 0.0050 | 0.0050 |
|  | 600 | tb | 0.1000 | 0.1000 | 0.1000 |
|  | 1800 | Fe | 0.0308 | 0.0098 | 0.095 |
|  | 1800 | E0 | 0.0050 | 0.0050 | 0.0050 |
|  | 1800 | tb | 0.1000 | 0.1000 | 0.1000 |
| *A. digitifera* | 10 | Fe | 0.002821705 | 0.0004314098 | 0.003223624 |
|  | 10 | E0 | 0.005 | 0.001 | 0.005 |
|  | 10 | tb | 10 | 7.726664 | 10 |
|  | 30 | Fe | 0.001332489 | 0.001135291 | 0.001492771 |
|  | 30 | E0 | 0.005 | 0.005 | 0.005 |
|  | 30 | tb | 10 | 8.721839 | 10 |
|  | 60 | Fe | 0.001898815 | 0.0008999647 | 0.002333828 |
|  | 60 | E0 | 0.005 | 0.001 | 0.005 |
|  | 60 | tb | 2.933751 | 2.133663 | 10 |
|  | 600 | Fe | 0.000365073 | 0.0002425592 | 0.00048346 |
|  | 600 | E0 | 0.005 | 0.005 | 0.005 |
|  | 600 | tb | 7.689085 | 4.539667 | 10 |
|  | 1800 | Fe | 0.0002340993 | 0.0001639761 | 0.000280067 |
|  | 1800 | E0 | 0.005 | 0.005 | 0.005 |
|  | 1800 | tb | 10 | 7.377308 | 10 |

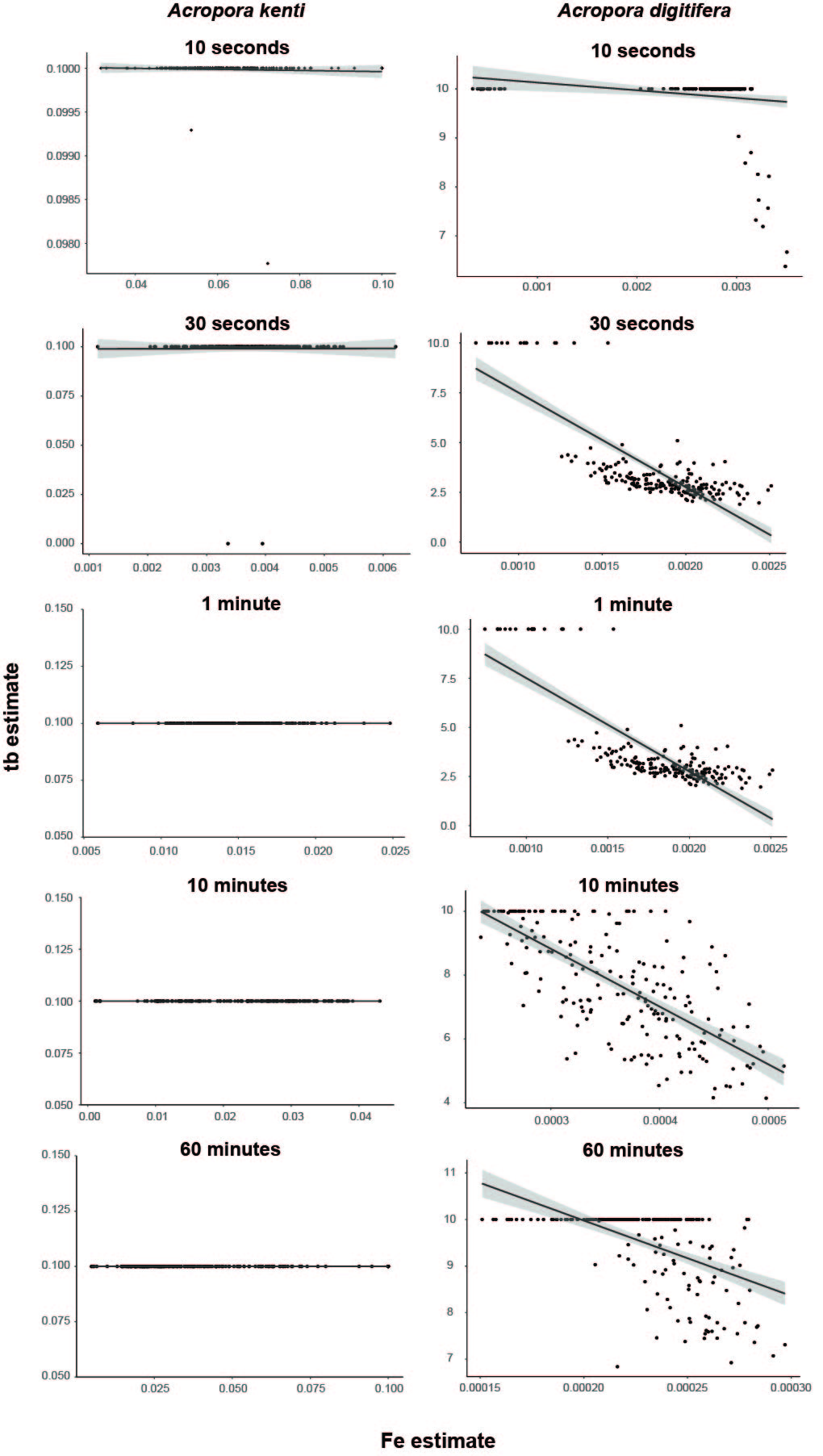

**Fig. S2** Correlation between the *Fe* and *tb* parameters across contact times following 200 bootstrap iterations of model optimisation. Points represent the value of each parameter estimate following each bootstrap run. Line and shaded region represent a linear smooth function and standard error used to explain the negative correlation

**Table S5** Spearman correlation values for *Fe* and *tb* parameters across contact time and species. Correlations are visualised in Figure S2.

| **Species** | **Contact time (s)** | **Spearman correlation** |
| --- | --- | --- |
| *A. kenti* | 10 | - 0.0214 |
|  | 30 | - 0.0052 |
|  | 60 | N/A |
|  | 600 | N/A |
|  | 1800 | N/A |
| *A. digitifera* | 10 | - 0.3834 |
|  | 30 | - 0.0208 |
|  | 60 | - 0.7214 |
|  | 600 | - 0.7112 |
|  | 1800 | - 0.5967 |

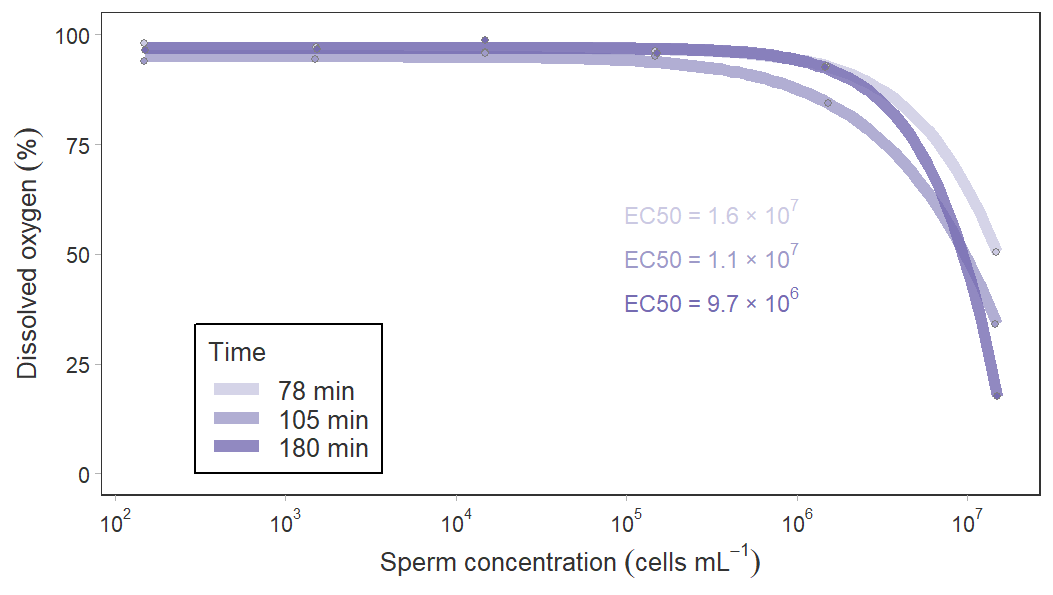

**Fig. S3** Relationship between sperm concentration and dissolved oxygen across three time-exposure gradients. Data points are provided at each sperm concentration. Each colour delineates each of the three exposure times and EC_50_ values for each exposure time is presented in the text with corresponding colour. Sperm concentration influenced dissolved oxygen (DO) at concentrations exceeding 10^5^ sperm mL^-1^, with DO approaching 50% or less at 10^7^ sperm mL^-1^. There was no measurable change in pH across the sperm concentrations, which ranged between 7.99 and 8.06
